# Translational impacts of enzymes that modify ribosomal RNA around the peptidyl transferase centre

**DOI:** 10.1101/2024.03.21.586085

**Authors:** Letian Bao, Josefine Liljeruhm, Rubén Crespo Blanco, Gerrit Brandis, Jaanus Remme, Anthony C. Forster

## Abstract

Large ribosomal RNAs (rRNAs) are modified heavily post-transcriptionally in functionally-important regions but, paradoxically, individual knockouts (KOs) of the modification enzymes have minimal impact on *Escherichia coli* growth. Furthermore, we recently constructed a strain with combined KOs of five modification enzymes (RluC, RlmKL, RlmN, RlmM and RluE) of the “critical region” of the peptidyl transferase center (PTC) in 23S rRNA that exhibited only a minor growth defect at 37°C (although major at 20°C). However, our combined KO of modification enzymes RluC and RlmE resulted in conditional lethality (at 20°C). Although the growth rates for both multiple-KO strains were characterized, the molecular explanations for such deficits remain unclear. Here, we pinpoint biochemical defects in these strains. *In vitro* fast kinetics at 20 and 37°C with ribosomes purified from both strains revealed, counterintuitively, the slowing of translocation, not peptide bond formation or peptidyl release. Rates of protein synthesis *in vivo*, as judged by the kinetics of β-galactosidase induction, were also slowed. For the five-KO strain, the biggest deficit at 37°C was in 70S ribosome assembly, as judged by a dominant 50S peak in ribosome sucrose gradient profiles at 5 mM Mg^2+^. Reconstitution of this 50S subunit from purified five-KO rRNA and ribosomal proteins supported a direct role in ribosome biogenesis of the PTC region modifications *per se*, rather than of the modification enzymes. These results clarify the importance and roles of the enigmatic rRNA modifications.

## Introduction

The ribosome is the complex ribonucleoprotein factory for cellular protein synthesis, termed 70S in *E. coli*. Initiation is carried out by the ribosomal small subunit, the 30S. It contains 16S rRNA that is modified posttranscriptionally, but the modifications have little effect on *in vitro* reconstitution and activity^[1]^. Both elongation and termination involve nucleophilic attacks catalyzed by the ribosomal large (50S) subunit. It contains unmodified 5S ribosomal RNA (rRNA) and 23S rRNA that is heavily modified, mostly with pseudouridylations and methylations^[2,3]^. Thirteen of these 25 modifications are clustered in domain V around the peptidyl transferase center (PTC) loop (Figure 1), suggesting functional importance (reviewed in ^[4,5]^). However, the impacts of rRNA modifications on ribosome biogenesis, assembly and function remain poorly understood.

**Figure 1.**
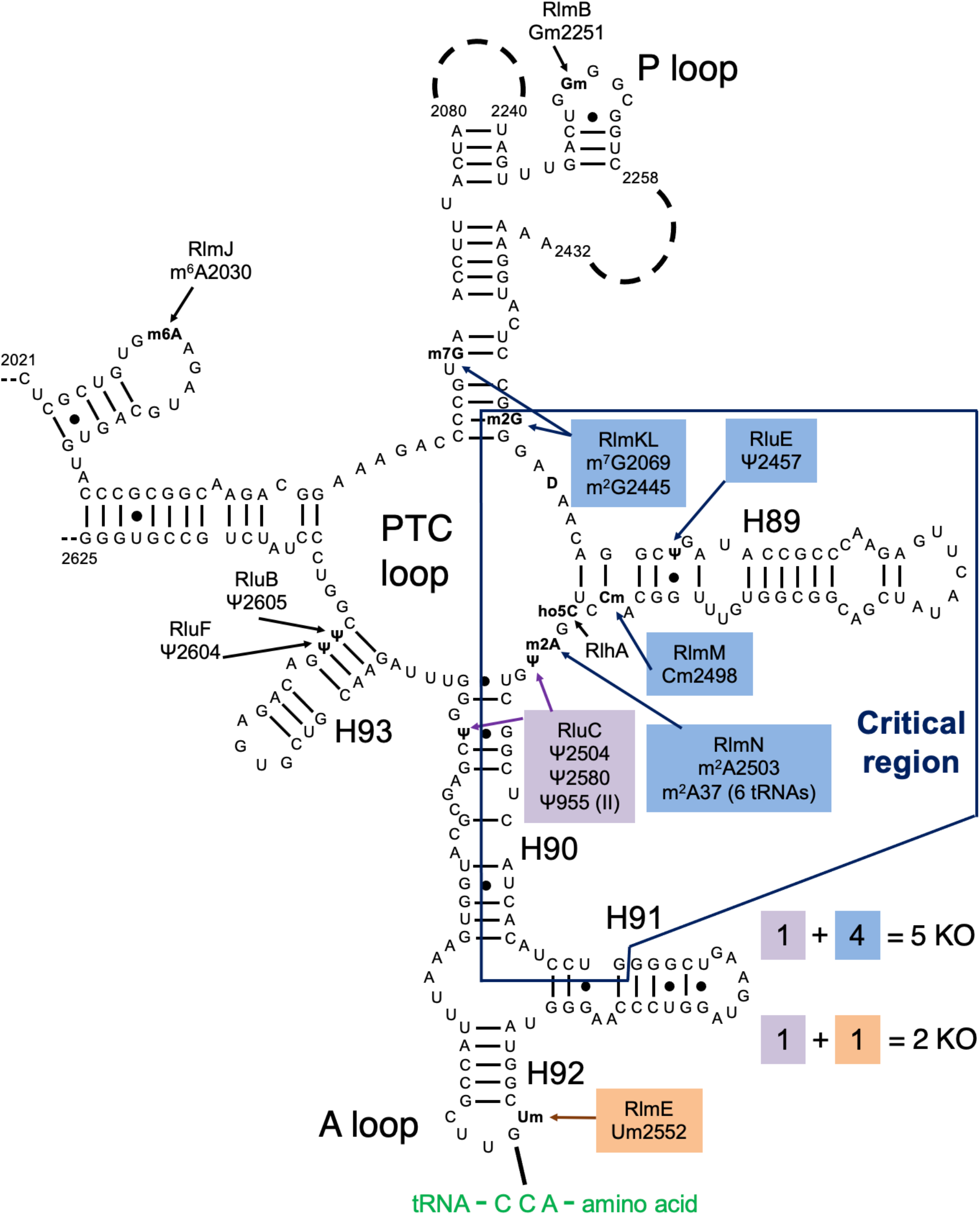
Our combination KOs of key rRNA modification enzymes for *E. coli*. The secondary structure contains most of domain V of the 23S rRNA, including the “critical region” (hollow box) around the PTC loop. Shown are all domain V modifications (ho5C is partial) and the modification enzyme KOs used in this study (filled boxes). Base pairing with aminoacyl-tRNA at the A site is shown. Modified from Liljeruhm *et al*. (2022) with permission.

Frustratingly, individual knockouts (KOs) of all rRNA modification enzymes resulted in very minimal growth deficits (e.g. ^[6,7]^). The notable exception, displaying a two-to fourfold decrease in growth rate, was *ΔrlmE* which abolished the 2’O-methylation of U2552 at the A site’s A loop adjacent to the PTC loop (Figure 1)^[8]^. An additional complication with KO studies is that non-modification “moonlighting” functions of the enzymes are possible. Indeed, *ΔrlmE* was partially rescued by overexpressing assembly factors EngA or ObgE^[9]^. But a catalytically-inert rlmE mutant did not rescue the *ΔrlmE* growth deficit^[10]^, implying a direct role for modification in growth. Moreover, depletion of SAM impacts ribosome biogenesis through hypomodification of Um2552^[11]^. Biochemically, *ΔrlmE* ribosomes exhibited deficits in 70S assembly^[9,12,13]^ and *in vitro* translocation^[14]^. The *ΔrlmE* strain accumulated ribosomal 45S intermediates with reduced levels of ribosomal protein^[8]^, including L5 and L19 that form intersubunit bridges between the 50S and 30S subunits^[15]^.

Due to the redundancy of individual bacterial rRNA modification enzymes, combination KOs are necessary to unmask their significance. In an *in vitro* study, Green and Noller^[16]^ reported a “critical natural element” of the 23S rRNA, an 80-nucleotide region around the PTC loop where modifications were essential for function *in vitro* (>100x stimulation; Figure 1). This was discovered by reconstituting 50S subunits with hybridized pairs of *in-vitro*-transcribed and isolated natural 23S rRNA portions and assaying for peptidyl transferase activity. This unphysiological assay, termed “the fragment reaction,” used a 5’-truncated fMet-tRNA^fMet^ fragment as the P-site substrate for the 50S, puromycin as the A-site substrate, and incubation in 33% methanol on ice. This study bears upon the *in vitro* synthesis of *E. coli* ribosomes (and ultimately self-replication)^[17]^, where the partially sequential, slow modification enzymology is one of the challenges (J Liljeruhm and AC Forster, unpubl.). Nevertheless, a crude cell-free system enables ribosome synthesis and evolution^[18]^.

An *E. coli* strain combining KOs of all 11 pseudouridines has been constructed, but the growth and ribosome biogenesis defects were minimal^[19]^. Recently, we reported the *ΔrluC/ΔrlmE* strain, which lacks psudouridylation of U955, U2504 and U2580 and methylation of U2552 (Figure 1)^[5]^. Happily, this strain displayed the most severe bacterial phenotype yet seen for rRNA modifications: a threefold growth deficit at 37°C and lethality at 20°C. Furthermore, its sucrose gradient profile revealed a severe decrease in the ratio of 70S to 50S. This is consistent with knowledge that ribosome assembly defects typically display cold sensitivity associated with incorrect rRNA folding^[20]^. We also constructed a second combination KO strain that lacked five modification enzymes, *ΔrluC/ΔrlmKL/ΔrlmN/ΔrluM/ΔrluE* (abbreviated *ΔCKLNMuE*; note that rlmE is not deleted), resulting in the absence of all modifications in the critical region except D and the partially-modified ho5C (Figure 1). Interestingly, it was viable and showed a minimal growth deficit at 37°^[5]^. However, at 20°C, it displayed a major growth deficit. Our prior kinetic analyses on these combined KO ribosomes were limited to tripeptide syntheses at 37°C for *ΔrlmE* and *ΔrluC/ΔrlmE*, where WT was 1.3x faster than *ΔrlmE* (consistent with ^[14]^) which was in turn 1.2x faster than *ΔrluC/ΔrlmE*^[5]^. Our other *in vitro* assays were less comprehensive, either limited to end timepoints of dipeptide synthesis at 37°C for *ΔrlmE* and *ΔrluC/ΔrlmE* with no differences detected compared to WT, or end timepoints of fragment reactions on ice in 33% methanol with *ΔCKLNMuE* 50S with a 2x decrease in product yield compared to WT. Furthermore, kinetics rates were untested at 20°C, the temperature where the most severe growth defects were measured, and translation rates *in vivo* at any temperature were unknown. Here, we further characterize these strains in order to determine their molecular defects.

## Materials and methods

### Materials

Tritium-labeled Met was purchased from PerkinElmer. All other chemicals and reagents were purchased from Sigma-Aldrich or Merck. *E. coli* 70S ribosomes, translation factors, and fMet-tRNA^fMet^ were prepared as described^[21]^. Preparation of the three chromosomal mutant strains in MG1655 has been detailed^[5]^. Note that MRE600 was used as “WT” for *in vitro* kinetics experiments (it contains lower RNase I activity when compared with MG1655 WT).

### Ribosome sucrose gradients

The sucrose gradient profile was recorded during large scale ribosome preparation as described^[21]^ with 5 mM Mg^2+^. Briefly, *ΔCKLNMuE* cells were grown to OD_600_ ∼0.8 at 37°C in LB medium and collected by centrifugation. The ultracentrifugation-based purification of ribosomes was performed as described^[22]^. The A_254_ of the sucrose gradient was read with optical unit UV-1 (Pharmacia Biotech) and recorded by BD-112 dual-channel chart recorder (Kipp & Zonen). All *ΔCKLNMuE* 50S subunits used here and in ^[5]^ were taken directly from the 50S fraction large peak of the sucrose gradient profile of total ribosomes. WT 50S subunits were purified from the 70S ribosome peak by decreasing Mg^2+^ to 3 mM to allow dissociation of the 50S and 30S, with the 50S fraction being collected by further sucrose gradient ultracentrifugation as for 70S purification.

### Fast kinetics *in vitro* by quench-flow

Synthetic XR7 mRNA was prepared as described^[23]^ with the sequence: 5′ GGGAAUUCGGGCCCUUGUUAACAAUU<u>AAGGAGG</u>UAUUAA **xxx xxx** UUGCAGAAAAAAAAAAA AAAAAAAAAA3′ The Shine–Dalgarno sequence is underlined and the coding sequences are in bold for fMet-Phe-Phe (AUG UUC UUC) or fMet-stop (AUG UAA). All kinetics experiments were conducted in HEPES-polymix buffer (pH 7.5) containing: 5 mM Mg(OAc)_2_, 95 mM KCl, 5 mM NH_4_Cl, 0.5 mM CaCl_2_, 1 mM spermidine, 8 mM putrescine, 1 mM dithioerythritol, and 30 mM HEPES-KOH pH 7.5. Active concentrations of ribosomes (typically ∼50%–75% of the total within the 70S peak on a sucrose gradient for all types of ribosomes here), fMet-tRNA^fMet^, EF-Tu and elongator tRNA^Phe^ were determined through the yield of dipeptide formed when the assayed component was limiting. For dipeptide formation, tripeptide formation and EF-G titration assays, 70S initiation complex (IC) and elongation mixture (EM) were formed by preincubating at 37°C for 20 min separately^[23]^. IC contained 2 μM 70S ribosomes, 5 μM XR7 fMFF mRNA, 2.4 μM f[^3^H]Met-tRNA^fMet^, 2 μM IF1, 2 μM IF2 and 2 μM IF3. For dipeptide formation, EM contained 5 μM EF-Tu, 5 μM tRNA^Phe^, 0.2 mM phenylalanine and 1 μM Phe-tRNA synthetase. For tripeptide formation, EM contained 20 μM EF-Tu, 5 μM EF-Ts, 5 μM EF-G (altered to 1 and 10 μM in EF-G titration assays), 10 μM tRNA^Phe^, 0.4 mM phenylalanine, and 2 μM Phe-tRNA synthetase. Both IC and EM were prepared in HEPES-polymix buffer (pH 7.5) supplemented with 1 mM ATP, 1 mM GTP, 10 mM phosphoenolpyruvate, 0.02 mg/mL pyruvate kinase and 0.005 mg/mL myokinase for energy regeneration. Fast *in vitro* kinetics assays were done at 20 and 37°C in a temperature-controlled quench-flow apparatus (RQF-3, KinTeck Corp.). Equal volumes of IC and EM were rapidly mixed, and the reactions were quenched with formic acid (17% final) at different time points. The samples were then centrifuged at 20,000g at 4°C for 30 min and the supernatant removed. Next, 120 μL of 0.5 M KOH was added to the pellet to hydrolyze all the peptides and unreacted f[^3^H]Met from the tRNAs at room temperature for 15 min. Formic acid was added (17% final) to precipitate the deacylated tRNAs. After centrifugation at 20,000g at 4°C for 30 min, the supernatant was analyzed by C18 reversed phase HPLC coupled with a β-RAM model 3 radioactivity detector (IN/US Systems). Separation of f[^3^H]Met-Phe-Phe, f[^3^H]Met-Phe and f[^3^H]Met was achieved by elution with 50% methanol/ 50% H_2_O/0.1% trifluoroacetic acid for 20 min at 0.45 mL/min. The fraction of [^3^H] peptide out of the total [^3^H] signal at each time point was calculated and the data fitted to a single exponential function with Origin 7.5 (OriginLab Corp.).

For the release assays, release complexes harboring f[^3^H]Met-tRNA^fMet^ in the P site and the UAA stop codon in the A site were prepared by incubation at 37°C for 20 min of IC mentioned above but with 5 μM XR7 fM-stop mRNA and 2 μM f[^3^H]Met-tRNA^fMet^ instead. This incubated IC was then diluted 10x with HEPES-polymix buffer (pH 7.5) and equilibrated at 37°C for 5 min. The factor mix containing 6 μM release factor 1 (RF1) in HEPES-polymix buffer (pH 7.5) was incubated at 37°C for 5 min. The rates of release were assayed based on *in vitro* quench-flow kinetics as for elongation assays both at 20 and 37°. The quenched samples were cooled on ice, and supernatants containing released f[^3^H]Met were separated from pellets containing the f[^3^H]Met-tRNA^fMet^ by centrifugation at 20,000g for 30 min at 4°C. Pellets were dissolved in 120 μL 0.5 M KOH at 37°C for 15 min and formic acid was added (17% final) to precipitate the deacylated tRNAs. The radioactive supernatants and dissolved pellets were diluted in ProFlow G+ scintillation liquid for counting (Beckman Coulter LS 6500). The fraction of release was calculated as:

Release fraction = [^3^H] supernatant/([^3^H] supernatant + [^3^H] pellet).

Kinetic rates were estimated by double exponential decay function fitting with Origin 7.5 (OriginLab Corp.). P values were calculated by one-tailed *t*-tests.

### *In vitro* reconstitutions of the 50S subunit

Extractions of 50S total proteins and rRNAs were as described^[24]^ but with ∼80 A_260_ units of 50S. The *in vitro* reconstitution was performed as described^[24]^ with minor changes. Briefly, 0.5 A_260_ units of total rRNAs was mixed with 0.6 equivalent units (e.u.) of 50S total proteins (TP50) in a 40 μL total volume containing 20 mM Tris-HCl (pH 7.5), 4 mM Mg(OAc)_2_, 0.4 M NH_4_Cl, 0.2 mM EDTA and 50 mM 2-meraptoethanol. 1 A_230_ unit of TP50 = 10 e.u., 1 A_260_ unit of 50S = 1 e.u. TP50. The mixture was then incubated at 44°C for 30 min. Then the Mg(OAc)_2_ concentration was increased to 20 mM and the solution incubated at 50°C for 90 min. This was then directly used without storage for fragment reactions (see below).

### *In vitro* peptidyl transferase reactions

Puromycin “fragment” reactions were performed as described^[16]^ with minor changes. Full-length f[^3^H]Met-tRNA^fMet^ (30 pmol) was used as the P-site substrate in a premix (total 60 μL) containing 50 mM Tris-HCl (pH 7.5), 400 mM KCl, 60 mM MgCl_2_, 1 mM puromycin and 15 pmol intact 50S subunit or 40 μL reconstituted 50S (see above). To this was added 30 μL cold methanol (= 33% final) for incubation on ice for 20 min. The reaction was stopped with 0.4 M final concentration of KOH and incubated at 37°C for 15 min. (This hydrolyzes any methanolysis side products; their yield is very minor and much lower than peptide product at all temperatures). Analysis was the same as for *in vitro* elongation assays (see above). The left 2 and right 2 bars in Sup. Figure 4A were replicated by a different person (results not shown).

### *In vivo* translation elongation rate assay

The *in vivo* translation rate was measured by β-galactosidase (LacZ) induction as described^[25]^. Overnight liquid cultures of WT MG1655 (in LB medium) and the KO strains (in LB+Km) were inoculated 1:100 in fresh LB medium (without Km). These were grown to OD_600_ ∼0.4 at 37°C and aliquoted into 500 μL/tube. The aliquots were pre-incubated at 20 or 37°C for at least 10 min. Then 5 mM IPTG was added to start the induction and, at desired time points, 208 μg of chloramphenicol was added to block further translation. “Quenched” samples were put on ice and the cells harvested by centrifugation at 20,000g for 1 min at 4 °C. ONPG-based colorimetric assays were performed exactly as described^[26]^ with 0.42 mg/mL chloramphenicol in Z buffer. Miller units were calculated and the LacZ induction time was plotted against square-root of lacZ activity (Schleif plot) to obtain the time for the first LacZ molecule synthesis. The X-axis intercept of the linear part of Schleif plot indicated the time (T_first_) for the ribosome to synthesize the first LacZ protein (1024 amino acids), therefore the elongation rate was calculated as 1024/T_first_.

### Constitutive exogenous protein expression assay

The exogenous reporter expression assays were performed closely to those described^[7]^. Their assays used plasmid pRFPCERtet with encoded β-lactamase wherein transformed cells were grown with or without anhydrotetracycline. In our assays, MG1655 and KO strains were transformed with pSB1C3 high-copy plasmids that express constitutively amilGFP, mRFP1 or fwYellow^[27]^. Transformed cells were grown in triplicates for 18 h at 37°C in 1 mL LB media with 33 μg/mL chloramphenicol (without Km). After incubation, cells were harvested by centrifugation at 20,000g for 1 min at 4°C and washed twice with PBS. The fluorescence of the cells was measured by an Infinite M200 pro plate reader (Tecan) at 485/535 nm for amilGPF, 531/595 nm for mRFP1 and 503/540nm for fwYellow. A_600_ was measured at the same time. Fluorescence per cell was calculated by fluorescence/A_600_. All values were normalized to WT. The KO ribosomes were not more sensitive than WT to chloramphenicol in the presence of chloramphenicol acetyltransferase encoded on the pSB1C3 vectors as judged by our protein expression yields.

### RlmE mutagenesis and rescue experiments

Catalytically-inert rlmE K38A^[10]^ was recreated by inverse PCR mutagenesis^[28]^ of RlmE in the low-copy pHB plasmid (derived from pSC101 with chloramphenicol resistance)^[29]^. Oligodeoxribonucleotide primers (Integrated DNA Technologies) had the sequences: 5’-GCTCTTGATGAAATACAGCAAAGTGAC-3’ (forward primer), and 5’-AAACCAGGCACGGGAAC-3’ (reverse primer). The plasmids were transformed into WT and *ΔrlmE* MG1655 strains. Bacterial growth was measured with a Spark® 10M Multimode Microplate Reader (Tecan) and Corning® 96-well flat-bottom plates (Sigma Aldrich).

## Results

### Effects of PTC modification enzyme KOs on translocation kinetics at different temperatures

The gaps in knowledge listed in the Introduction concerning the elongation kinetics of our combined KO strains^[5]^ were now filled (Sup. Figure 1; summarized in Sup. Table 1 and Figures 2A-D). Dipeptide syntheses were about 4x slower at 20°C (Figure 2A) than 37°C (Figure 2B) with the different ribosomes giving very similar results (Sup. Table 1). Thus, despite the above-mentioned defect for *ΔCKLNMuE* 50S in the fragment reaction on ice in 33% methanol^[5]^, peptide bond formation rates at 20 and 37°C in physiological buffer were unaffected in all the KOs and could not explain cold-sensitive bacterial growth. Thus, this defect versus WT in 33% methanol on ice^[5]^ seems irrelevant to the physiological situation (supported also by Sup. Figure 4A discussed below).

**Figure 2.**
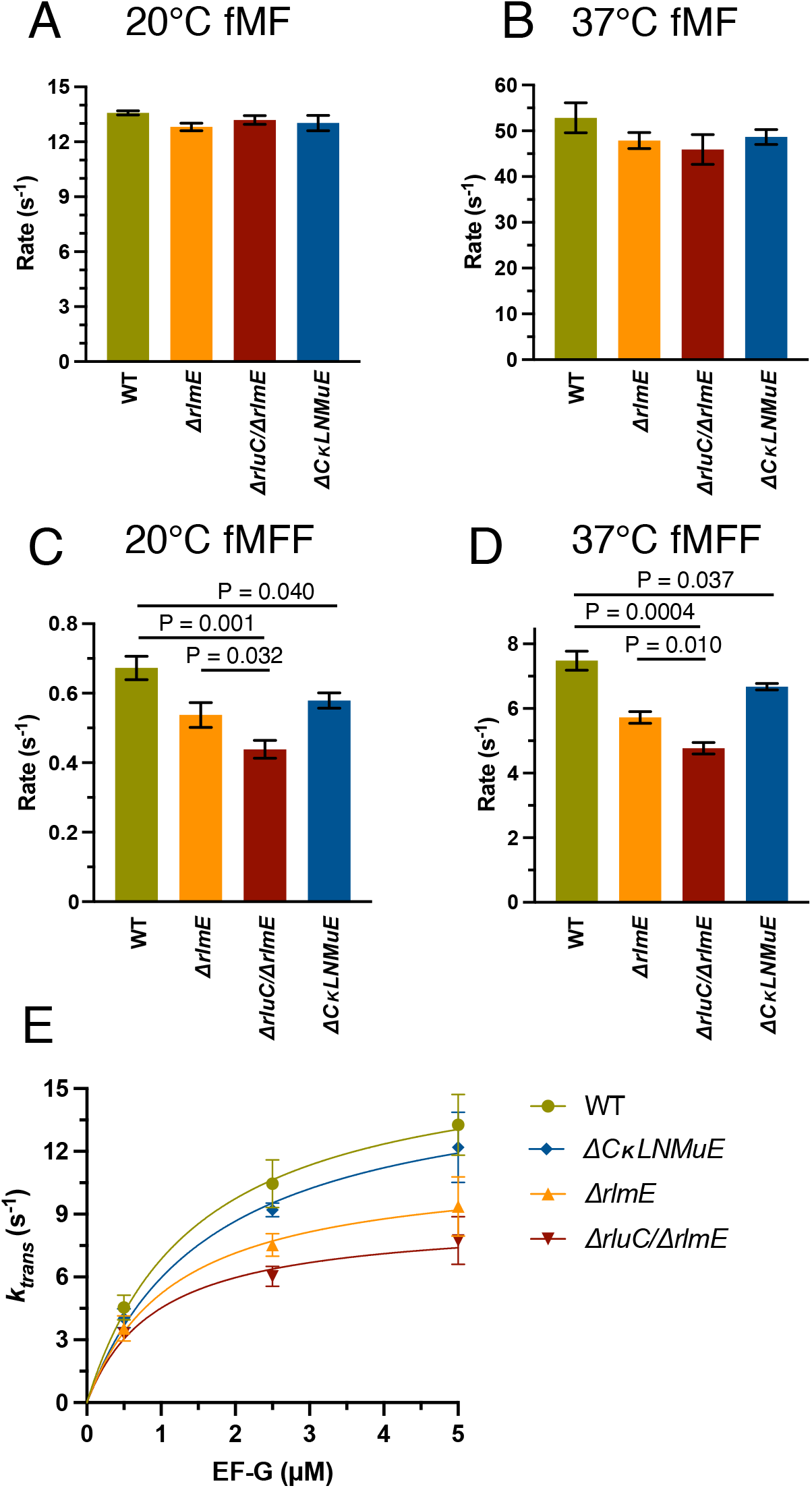
*In vitro* fast kinetics-based elongation assays of KO ribosomes. fMet-Phe dipeptide formation rates at 20°C (A) and 37°C (B), and fMet-Phe-Phe tripeptide formation rates at 20°C (C) and 37°C (D). (E) Calculated translocation rates at 37°C with different EF-G concentrations. Error bars are standard errors, n ≥ 2. P values are not shown for non-significant comparisons. Left three bars in Fig. 2D taken from Liljeruhm *et al*. (2022).

Having found significant reductions in tripeptide synthesis rates of the *ΔrluC/ΔrlmE* ribosomes at 37°^[5]^ that correlated with the major growth deficit of the strain at that temperature, we now tested *ΔCKLNMuE* ribosomes (Figure 2D, blue bar). Indeed, tripeptide synthesis by *ΔCKLNMuE* ribosomes at 37°C was reduced, despite the mild growth defect at that temperature *in vivo*. Given the more severe phenotype of *ΔCKLNMuE* at colder temperature^[5]^, we further tested tripeptide synthesis with *ΔCKLNMuE* ribosomes at 20°C (Figure 2C, blue bar) and again found a defect. The same assays were applied to *ΔrlmE* and *ΔrluC/ΔrlmE* ribosomes (Figure 2C, orange and red bars). Interestingly, although *ΔrluC/ΔrlmE* is lethal at 20°C^[5]^, its 70S ribosomes harvested at 37°C were still able to catalyze tripeptide formation at 20°C at 69% of the WT rate. Compared to *ΔrlmE*, which is viable at 20°C^[5]^, the tripeptide synthesis rate for *ΔrluC/ΔrlmE* was significantly less. These results indicated that slow tripeptide, not dipeptide, synthesis contributes to the cold-sensitivities of all of the KO strains.

To pinpoint whether the binding of translocation factor EF-G or a subsequent translocation step was affected, EF-G was titrated at 37°C and plotted against actual translocation rates calculated as

*translocation time = tripeptide synthesis time – 2x dipeptide synthesis time*

(Figure 2E and Sup. Table 2). For the ribosomes of all strains, curves, rather than straight lines, were obtained that indicated near-saturation at high EF-G concentration (5 μM = 2x standard concentration) and not weaker EF-G binding by the *ΔrlmE* and *ΔrluC/ΔrlmE* ribosomes (based on similar EF-G k_cat_/K_M_ values listed in Sup. Table 2 legend). This suggested that peptidyl-tRNA movement on the ribosome, rather than just weaker EF-G binding, was the affected translocation step (see right graphical image).

Because it cannot be ruled out that the defects are due to possible moonlighting functions of the modification enzymes rather than the absence of rRNA modifications *per se* (see Introduction), we performed two types of experiments here that bear upon this. Firstly, we confirmed the control of ^[10]^ by reconstructing their catalytically-inert RlmE K38A mutant and showing that it could not rescue the growth defect of *ΔrlmE* (Sup. Figure 2). Furthermore, we found that expression of the mutant even inhibited WT and *ΔrlmE* growth (Sup. Figure 2; RlmE was selected for this control because it’s KO gave the largest growth inhibition of the single modification enzyme KOs^[6,7]^). Secondly, we performed five-KO ribosome reconstitutions *in vitro* where the only difference between the tests and controls was the absence of the modifications (not the modification enzymes), yet we still detected catalytic defects (see Figure 4C below). These experiments (^[10]^, Sup. Figure 2 and Figure 4C) support direct roles of the rRNA modifications *per se* in the phenotypes^[5]^ of the KO strains.

### Translation termination at different temperatures is unaffected

In addition to peptide bond formation, the PTC catalyzes one other chemical reaction: hydrolysis of the polypeptide from the peptidyl-tRNA in translation termination. We thus wondered whether or not the modification enzyme deficits would affect a standard release assay (Figure 3A). Release rates were about 1.3x slower at 20°C (Sup. Figure 3A and Figure 3B) than 37°C (Sup. Figure 3B and Figure 3C) but the different ribosomes gave similar results at a particular temperature. *ΔrluC/ΔrlmE* and *ΔCKLNMuE* displayed slightly faster but insignificantly different release rates compared to WT at 20°C (summarized in Sup. Table 1). This indicated that effects on release did not cause the phenotypes^[5]^ of the KO strains. Taken together, the unaffected dipeptide bond formation and release rates by the modification enzyme KOs measured here imply that the KOs do not substantially affect the PTC-catalyzed chemical reactions of the ribosomes around physiological-type conditions.

**Figure 3.**
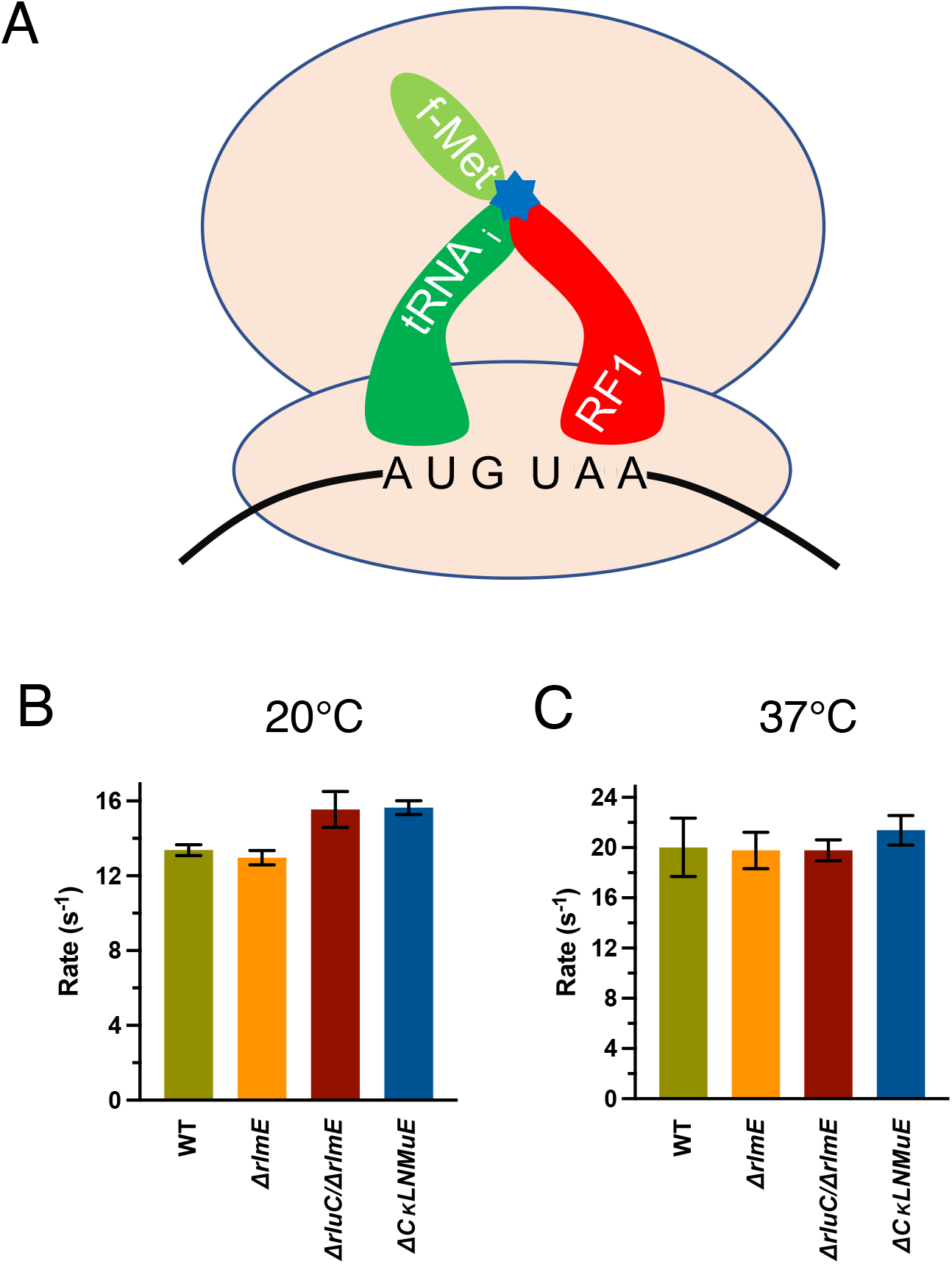
*In vitro* fast kinetics-based release assays of KO ribosomes. (A) Drawing of release assay depicting RF1-catalyzed hydrolysis (blue star) of the fMet acyl linkage on the initiator tRNA. Rates at 20°C (B) and 37°C (C). Error bars are standard errors, n ≥ 2. Calculated P values > 0.05 based on one-tailed *t*-test.

### Effects on ribosome folding and assembly

Our prior sucrose gradient profiling of ribosomes^[5]^ found a major defect in 70S assembly for *ΔrluC/ΔrlmE* but not *ΔCKLNMuE*, despite the latter lacking most of the “critical region” modifications. For *ΔCKLNMuE* ribosomes, assembly was only slightly sensitive to the cold. However, these conclusions were based on analytical profiles performed using the standard Mg^2+^ concentration of 10 mM, which is unphysiologically high and stabilizing. When we did large-scale purifications of the 70S ribosomes at 5 mM Mg^2+^, which is closer to the estimated physiological concentration of free Mg^2+^ (1-2 mM)^[30]^, the ratio of 70S to 50S ribosomal particles decreased greatly for *ΔCKLNMuE* compared to the WT (Figure 4A). This indicates that defective assembly of the 70S ribosome is a major contributor to the phenotype of the *ΔCKLNMuE* strain.

**Figure 4.**
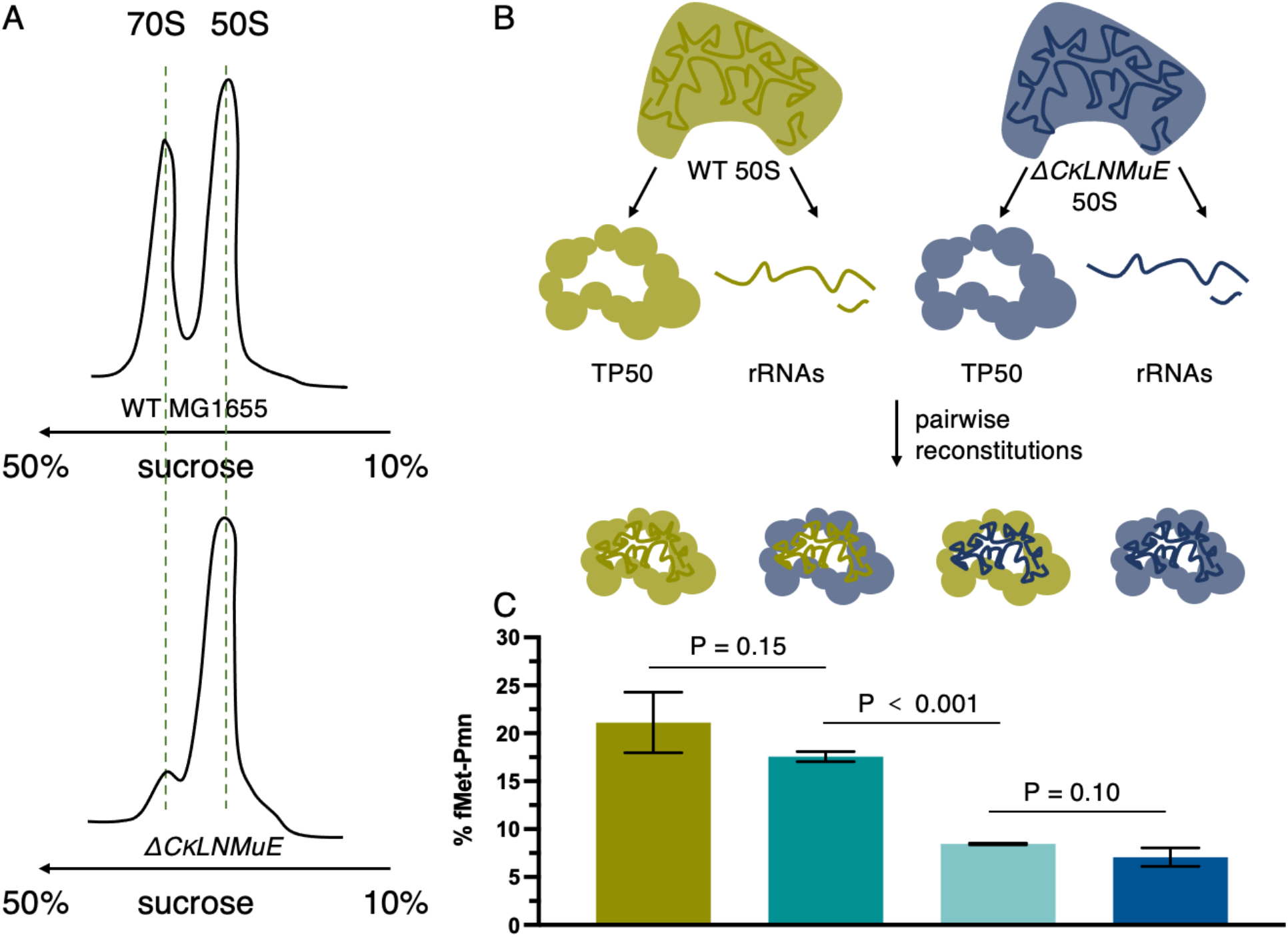
Ribosome sucrose gradient profiles and *in vitro* reconstitution assays with *ΔCKLNMuE*. (A) Representative ribosome sucrose gradient profiles at 5mM Mg^2+^ of *ΔCKLNMuE* and WT ribosomes. (B) Drawing of extractions of total 50S rRNAs and total 50S proteins and their reconstitutions. (C) Fragment reactions at 37°C of respective reconstitutions drawn directly above. Error bars are standard errors, n = 4.

Regarding ribosome assembly without most or all of the critical region modifications, we wondered if our much smaller defect (twofold) in the fragment reaction measured after folding *in vivo*^[5]^ compared with the huge defect after folding *in vitro* (>100x in ^[16]^) was due to *in vivo* versus *in vitro* folding conditions. Armed with sufficient purified 50S subunits from the WT and *ΔCKLNMuE* strains, we prepared total rRNAs and total proteins (TP50) from them for *in vitro* reconstitutions (Figure 4B). Firstly, we titrated the rRNA with TP50 in the standard two-step reconstitution to find that the optimized ratio was 1.0 A_260_ unit of rRNA to 1.2 equivalent units (e.u.) of TP50 (see Methods). The standard fragment reaction (on ice) was used to assay the activity of the reconstituted ribosomes, as well as reactions at 20 and 37°C to better model physiological conditions. The latter reactions were possible because we found that fragment reactions at 20 and 37°C were more efficient than those on ice (Sup. Figure 4A; note that reactions were very slow without methanol (not shown)). At 37°, the reconstituted *ΔCKLNMuE* 50S showed a threefold decrease in activity compared to reconstituted WT 50S (Figure 4C, first and last bars) which in turn had half of the activity of purified WT 50S (not shown). To rule out the possibility that the total proteins, rather than the undermodified rRNAs, caused the deficit in activity, we mixed the WT rRNAs with *ΔCKLNMuE* TP50 and *vice versa* (Figure 4C, middle two bars). Indeed, the activities tracked with the rRNAs, not the proteins. Similar results were obtained on ice (Sup. Figure 4B). Thus, the major differences between our results and those of ^[16]^ cannot be attributed to *in vivo* versus *in vitro* folding and presumably reflect other differences between the experiments (e.g. different combinations of modifications or their hybridization of non-covalently-linked 23S rRNA pieces). Also, our threefold deficits can now be attributed directly to the lack of PTC modifications rather than the lack of modification enzymes.

### Effects on elongation rates of endogenous protein synthesis *in vivo*

Given the slower tripeptide synthesis rates *in vitro* for KO ribosomes (^[5,14]^ and here), we wondered how these correlated with *in vivo* protein synthesis. Therefore, we measured protein synthesis elongation rates *in vivo* using the standard β-galactosidase (lacZ) induction assay^[25]^ at 20 and 37°. WT and KO strains were grown at 37°C and IPTG was added at the desired temperatures to induce synthesis of β-galactosidase. Our WT elongation rate of 15 amino acids per second at 37°C (Figure 5B) was comparable to published data^[31]^. At 20°C, WT displayed a 1.5x faster elongation rate than *ΔrlmE* and *ΔCKLNMuE* (Figure 5A and Sup. Figure 5A), correlating with their relative growth rates at that temperature^[5]^. Notably, *ΔrluC/ΔrlmE* showed a 4.5x slower rate than WT, and 3x slower rate than *ΔrlmE* and *ΔCKLNMuE*, taking ∼14 min to synthesize the first β-galactosidase compared with 3.6 min for WT (Figure 5A and Sup. Figure 5A). Such slow protein synthesis potentially causes the lethality of the *ΔrluC/ΔrlmE* strain at 20°C (see Discussion). At 37°, *ΔrlmE* and *ΔrluC/ΔrlmE* showed significantly slower rates than WT, but *ΔCKLNMuE* displayed a similar rate to WT (Figure 5B and Sup. Figure 5B). The data correlated well with the *in vitro* tripeptide synthesis rates (^[5,14]^ and Figure 2D) and the minimal deficit in growth rate for *ΔCKLNMuE* at 37°^[5]^.

**Figure 5.**
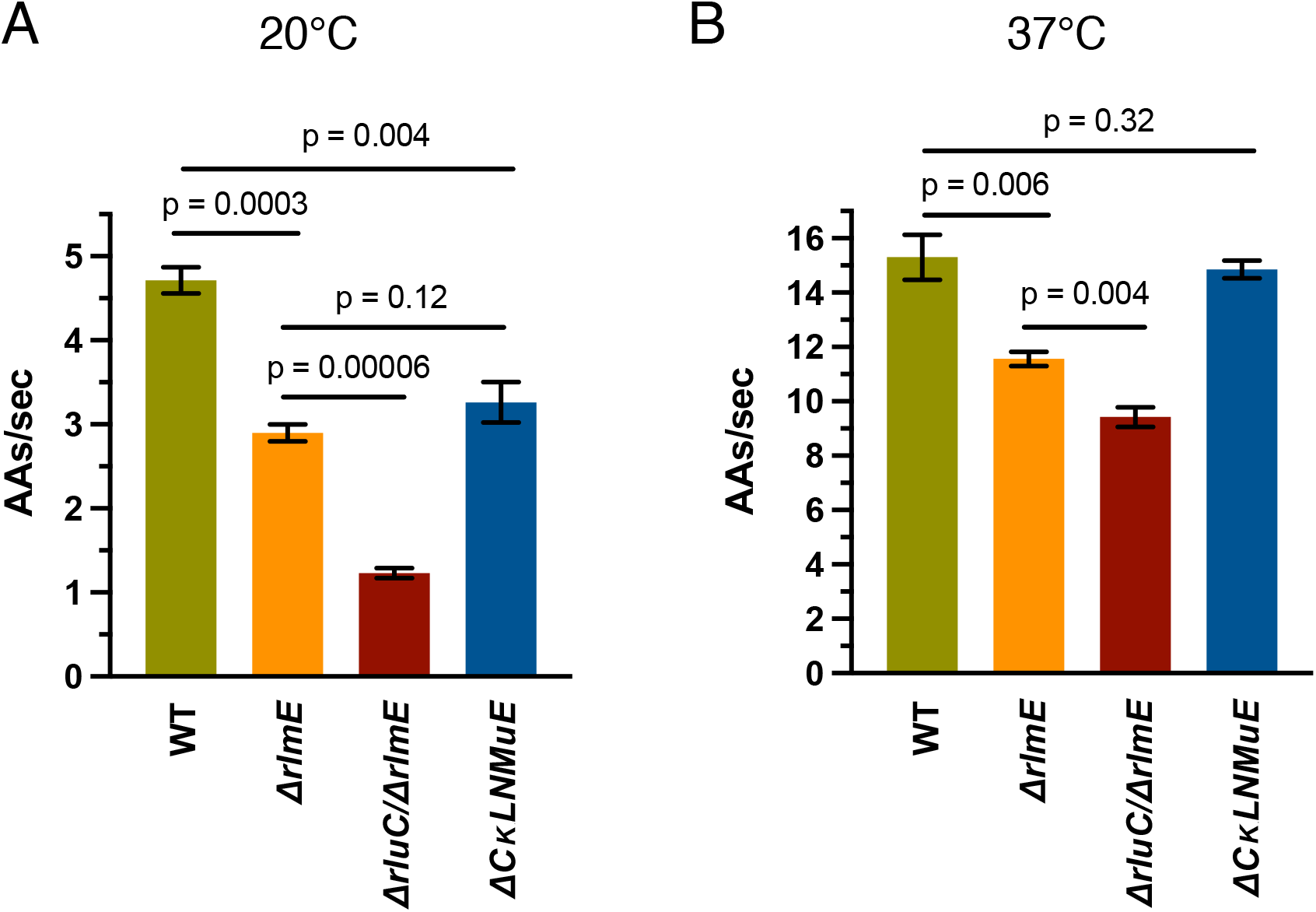
Influence of rRNA modification enzyme KOs on *in vivo* elongation rates measured by β-galactosidase induction at 20°C (A) and 37°C (B). Error bars are standard errors, n = 3.

### Effects on bacterial growth rather than exogenous protein overexpression

Recently, Pletnev *et al*.^[7]^ reported an *in vivo* protein overexpression assay that found surprisingly significant inhibitions even for most single KOs of *E. coli* rRNA methylation enzymes despite unaffected bacterial growth. Their method was based on overexpression of two fluorescent proteins (FPs) from the GFP family. We therefore tested this method with our KO strains using three related fluorescent proteins, amilGPF, mRFP1 and fwYellow, chosen because their overexpression in *E. coli* occurs without inclusion bodies or high toxicity^[27]^.

Plasmids constitutively expressing the FPs were transformed into *ΔrluC/ΔrlmE* and *ΔCKLNMuE* competent cells and the OD_600_ (Sup. Figure 6A) and fluorescence values were measured after 18h incubation at 37°C as published^[7]^. Unexpectedly, FP expression in the KO strains was not reduced compared to WT as judged by fluorescence/OD_600_ (Sup. Figure 6C) or SDS-PAGE which includes measurement of any unfolded FPs (Sup. Figure 6D). Also, the growth rates of *ΔrluC/ΔrlmE* and *ΔCKLNMuE* were suppressed somewhat by overexpression (e.g. for mRFP1, see Sup. Figure 7). Our unexpected finding of reductions in growth rates rather than FP overexpression may reflect unanticipated technical difficulties of this new assay. Also, it makes sense to rely more on the β-galactosidase assay (see above) for *in vivo* measurements because it is well-established, has been verified in comparison with other methods (e.g. see Figure S3J of ^[31]^) and it is more physiological than overexpression.

## Discussion

Following up on the unexpected growth phenotypes of both of our combination KOs of enzymes that modify around the PTC^[5]^, the data here provide molecular explanations. For both *ΔrlmE* and *ΔCKLNMuE*, evidence is provided that the growth defects were due to the actual absence of the rRNA modifications, rather than just the absence of the modification enzymes (Figure 4C and Sup. Figure 2). As previously noted for *ΔrlmE* ^[9,12-14]^ and *ΔrluC/ΔrlmE*^[5]^, a decrease in translocation rate *in vitro* (Figure 2D) and a severe defect in 70S assembly (Figure 4A) is also identified here for *ΔCKLNMuE. In vitro* defects in translocation, not dipeptide synthesis or release, are also demonstrated here at the temperature (20°C, Figure 2C) that caused the most severe growth inhibitions for *ΔrluC/ΔrlmE* and *ΔCKLNMuE*^[5]^. But comparison of the 20°C data with that at 37°C (Figures 2C, D) did not reveal cold sensitivity *in vitro* that could account for the cold sensitivity *in vivo*. However, defects in protein synthesis *in vivo* are now identified for the KO strains (Figure 5) that correlate with the growth defects at different temperatures^[5]^.

In theory, the % increase in time for a single elongation cycle by the KO ribosomes *in vitro* is expected to cause the same % increase in time for synthesis of the full-length β-galactosidase protein *in vivo* at the same temperature. This is assuming that the measured delay time in synthesis of the 1024-amino-acid β-galactosidase is largely due to elongation, not initiation, which is apparently true for WT (see Figure S3J of ^[31]^). Indeed, there was excellent agreement between the calculated pairs of *in vitro* and *in vivo* elongation cycle times for the three mutants at 37°C (Sup. Table 4). But at 20°C, our *in vivo* times were longer than expected compared with *in vitro* (Sup. Table 4), although the trend compared with WT was the same as at 37°. Perhaps *in vivo* the translocation defects are potentiated more at 20°C than at 37°C by the major 70S assembly defects associated with the three mutant strains^[5]^ and Figure 4A); this would shift rate limitation in the *in vivo* assay more to initiation (mindful that the *in vitro* values calculated in Sup. Table 4 only include elongation cycle steps). Regardless of the particular steps of protein synthesis affected at 20°C, our *in vivo*-measured large drop from 2.9 amino acids/sec for *ΔrlmE* to 1.2 amino acids/sec for *ΔrluC/ΔrlmE* (Figure 5A) could be responsible for the conditional lethal phenotype of the latter. This conclusion is based on the known correlation between elongation and growth rates at different temperatures and the extremely slow doubling time extrapolated from around 1.2 amino acids/sec on Figure 3 in ^[32]^.

It is puzzling that mature 70S ribosomes from our strains with combination KOs of enzymes that modify around the PTC both exhibited a defect, not in the two PTC functions of peptide bond formation and release, but rather in translocation. Similar kinetics results were published by others for *ΔrlmE*^[14]^. The ratcheting movements between the subunits that occur during translocation are distal from the PTC. However, a structural study of mature 50S particles from the *ΔrlmE* strain revealed that the largest rRNA displacements were of U2554 and G2553^[14]^, the latter being the A-site nucleotide which base pairs with the CCA end of aminoacyl-tRNA (immediately adjacent to Um2552 as shown in Figure 1) during translation^[33]^. After peptide bond formation occurs, the translocation steps include breaking this base pair. But dipeptide formation requires formation of this base pair and it is unknown if the U2554 and G2553 displacements occur in mature 70S from *ΔrlmE*. Another consideration is that, in contrast to translocation, both dipeptide formation and release are single-round nucleophilic attacks catalyzed by the PTC without as large a conformational change of the ribosome as occurs in translocation ratcheting^[34,35]^, so translocation may be more sensitive to structural changes.

Also hard to explain is defective 70S formation in all three KO strains, given that the PTC region is distal from the 50S subunit interface (see left graphical image). However, an assembly deficit seems reasonable based on the ribosomal literature (e.g. ^[14,20,36]^). Ribosome assembly defects typically display cold sensitivity^[20]^, *ΔrlmE* cells that lack a modification near the PTC accumulate pre50S particles with rRNA structural differences and ribosomal protein absences that cluster near the PTC^[14]^, and PTC formation is the final step in 50S maturation necessary for binding to the 30S^[36]^. After PTC maturation, no further conformational change in the 50S is required for 70S formation because the 50S has the same 3D structure whether or not it is bound to the 30S^[37]^. Although our 5KO 50S preparation that failed to form 70S functions quite well in fragment reactions here (with or without prior reconstitution), this can be rationalized by the use of unphysiologically-high Mg^2+^ concentrations which are known to facilitate PTC maturation. For *E. coli* 50S reconstitutions *in vitro*, a heat activation step at unphysiologically-high Mg^2+^ concentration is needed to fold the disordered rRNA within the peri-PTC region of helices H90-H93 (Figure 1)^[38]^.

In summary, different KO combinations around the PTC, though differing in phenotypic severity, exhibited remarkably similar molecular defects in translation. Evidence indicates that the enigmatic PTC region modifications act in synergy to correctly fold domain V of the 50S to allow efficient 70S assembly and translocation.

## Acknowledgements

We are grateful to Nicola Freyer for pilot fragment reactions, Raymond Fowler for purified translation factors and technical assistance, and Leif Kirsebom and Michael Pavlov for discussions. This work was supported by the Carl Tryggers Foundation, the Tore Nilsons Foundation and the Åke Wibergs Foundation (CTS22:1886, 2022-004 and M22-0037 to G.B.), the Estonian Ministry of Education and Research (PUT PRG1179 to J.R.) and the Swedish Research Council (NT project grant 2017-04148 to A.C.F.).

## Author Contributions

LB, JR & ACF conceptualized the study; LB, JL, RCB and GB obtained the data; all authors analyzed the data; LB & AF wrote the manuscript and all authors edited it.

## Data availability

The data supporting the study are presented in the figures and tables in the main text and the supplementary material and are available in more detail upon request from LB and AF.

---

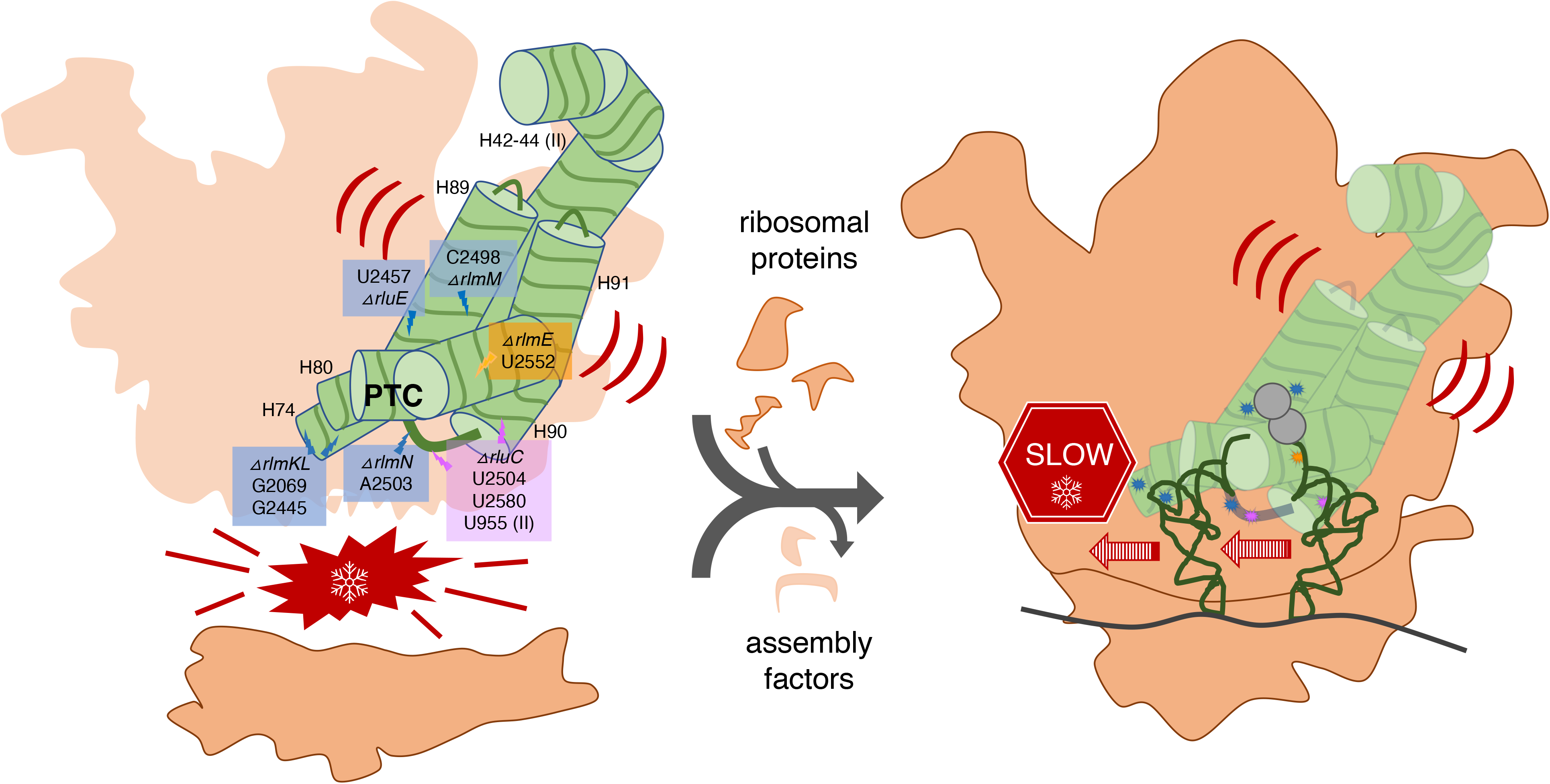

