## Supplemental Tables 1-4, Figures 1-7 for "Translational impacts of enzymes that modify ribosomal RNA around the peptidyl transferase centre"

**Translational effects of enzymes that modify**

**around the peptidyl transferase centre**

Contents

**Table S1.** Summary of translation and release rates of KO ribosomes at 20 and 37℃ for Figs. 2A-D & 3.

**Table S2.** Summary of calculated translocation rates of KO ribosomes at 37℃ for Fig. 2E.

**Table S3.** Summary of *in vivo* β-galactosidase synthesis rates of KO ribosomes at 20 and 37℃ for Fig. 5A and B.

**Table S4.** Comparison of fold changes (normalized to WT) of times for a single elongation cycle *in vitro* (calculated as τ*_fMFF_* – τ*_fMF_* with 2.5 μM EF-G from Sup. Table 1) and *in vivo* (calculated as 1/(amino acids/sec) from Fig. 5).

**Figure S1.** *In vitro* fast kinetics-based elongation assays of KO ribosomes.

**Figure S2.** Rescue experiments by expressing functional or catalytically-inert RlmE enzymes in WT or *ΔrlmE* strains.

**Figure S3.** *In vitro* fast kinetics-based release assays of KO ribosomes.

**Figure S4.** Comparison of fragment reactions at different temperatures.

**Figure S5.** Time courses of β-galactosidase induction *in vivo* (Schleif *et al.*, 1973) at 20℃ (A) and 37℃ (B).

**Figure S6.** Influence of rRNA modification enzyme KOs on overexpression of fluorescent proteins and on growth.

**Figure S7.** Influence of constitutive overexpression of mRFP1 on growth of *ΔCKLNMuE* and *ΔrlmE/Δrlu*C.

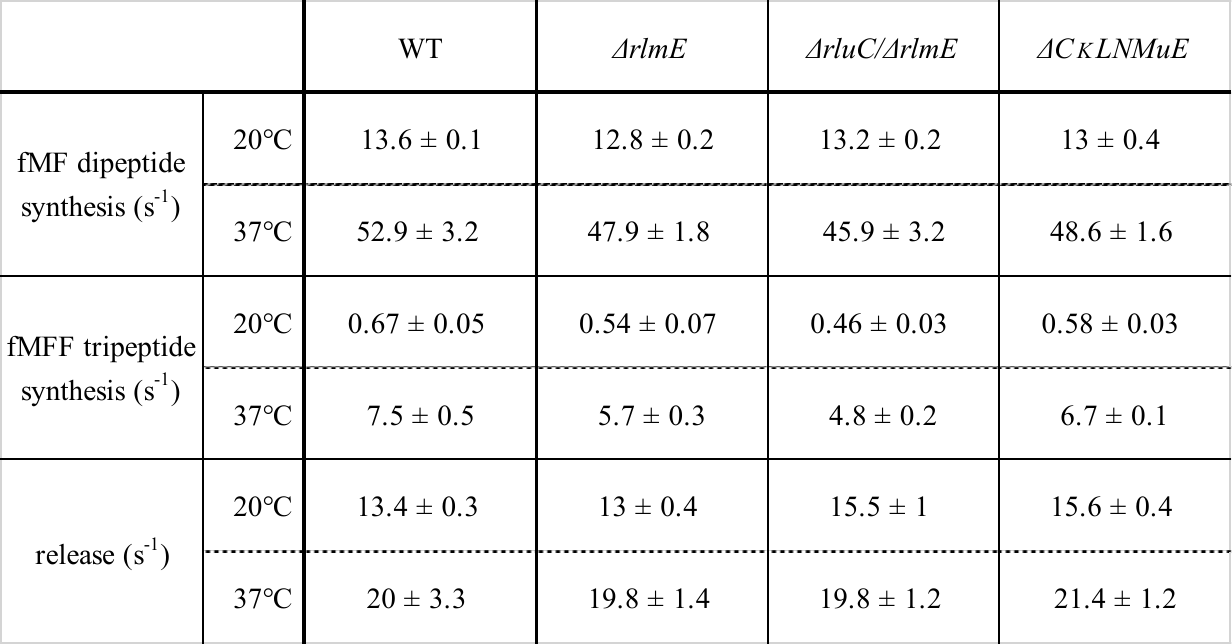

**Sup. Table 1.** Summary of translation and release rates of KO ribosomes at 20 and 37℃ for Figs. 2A-D & 3.

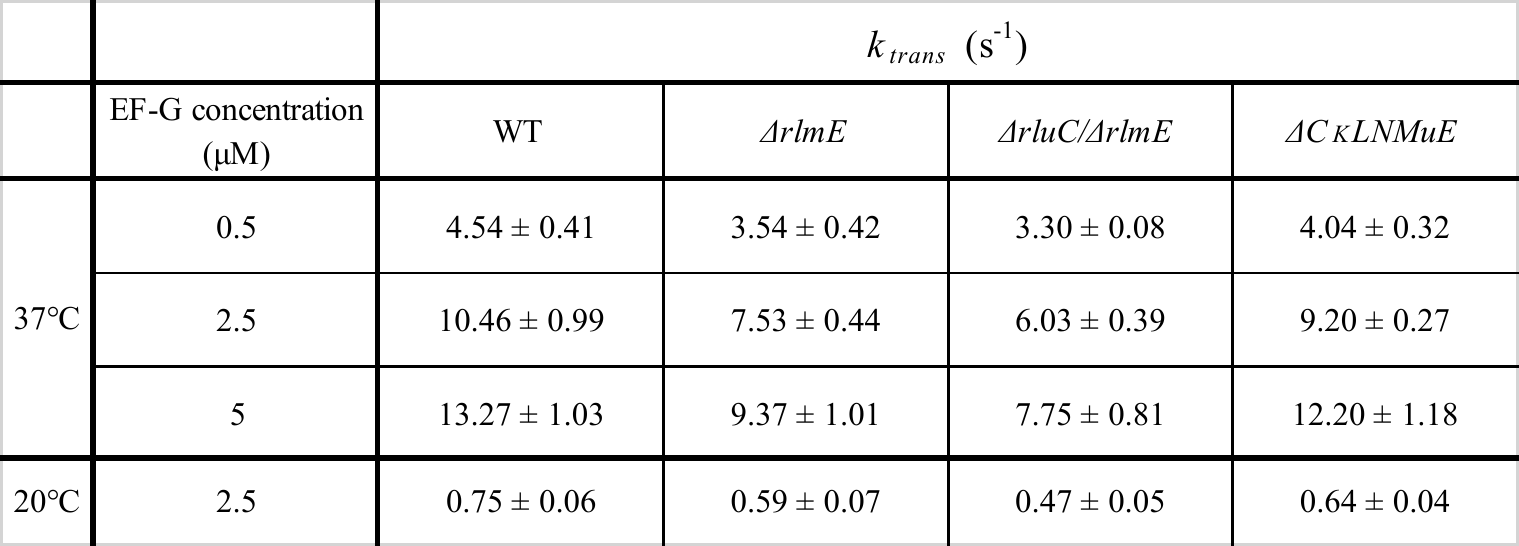

**Sup. Table 2.** Summary of calculated translocation rates of KO ribosomes at 37℃ for Fig. 2E. Values at 20℃ are included for comparison. Calculated k_cat_/K_m_ values at 37℃: WT, 11.5; *ΔrlmE*, 9.6; *ΔrluC/ΔrlmE*, 9.3 and *ΔCKLNMuE*, 9.5.

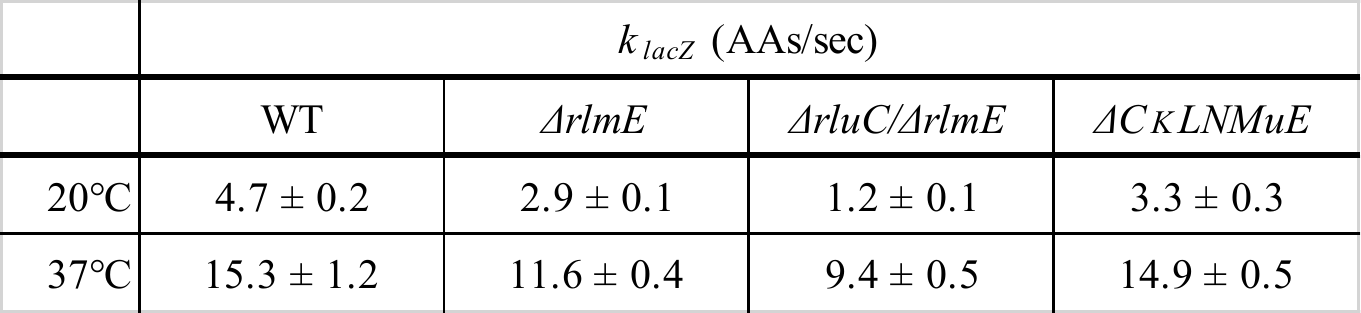

**Sup. Table 3.** Summary of *in vivo* β-galactosidase synthesis rates of KO ribosomes at 20 and 37℃ for Fig. 5A and B.

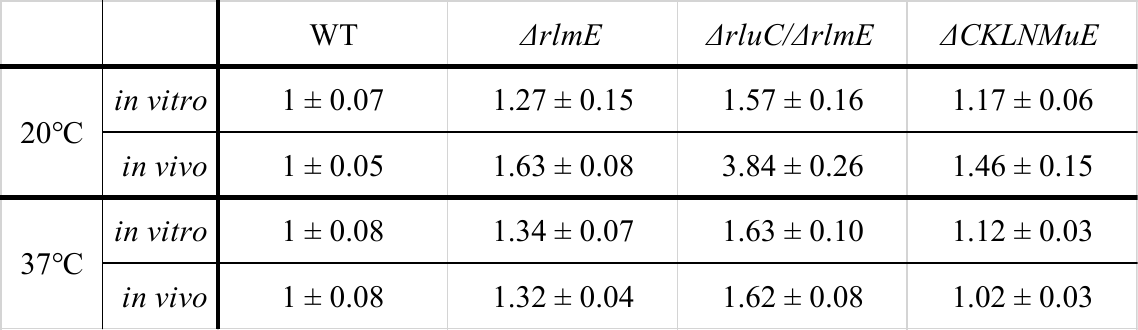

**Sup. Table 4.** Comparison of fold changes (normalized to WT) of times for a single elongation cycle *in vitro* (calculated as τ*_fMFF_* – τ*_fMF_* with 2.5 μM EF-G from Sup. Table 1) and *in vivo* (calculated as 1/(amino acids/sec) from Fig. 5).

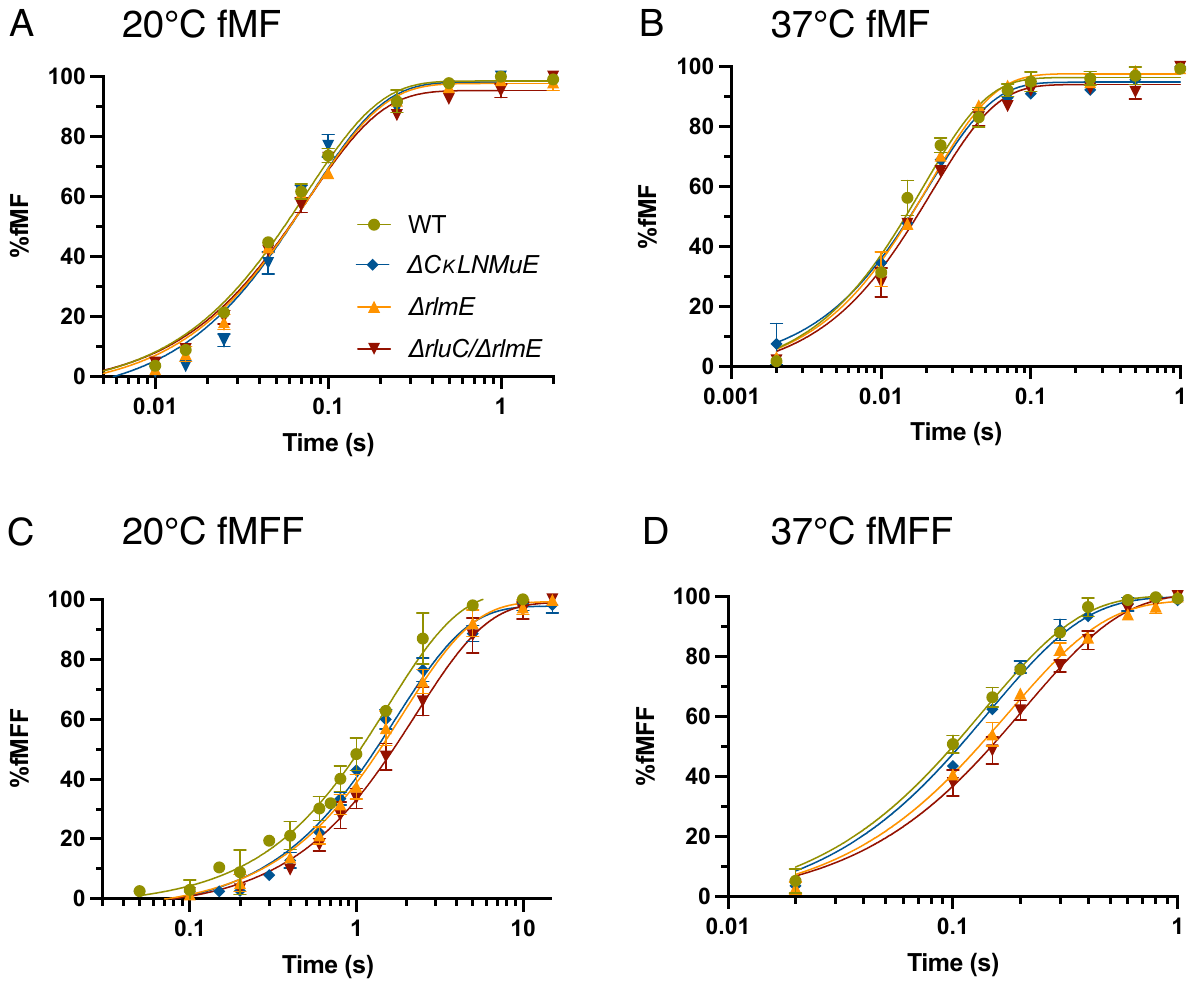

**Sup. Figure 1.** *In vitro* fast kinetics-based elongation assays of KO ribosomes. fMet-Phe dipeptide formation time courses at 20℃ (A) and 37℃ (B), and fMet-Phe-Phe tripeptide formation time courses at 20℃ (C) and 37℃ (D). Error bars are standard errors.

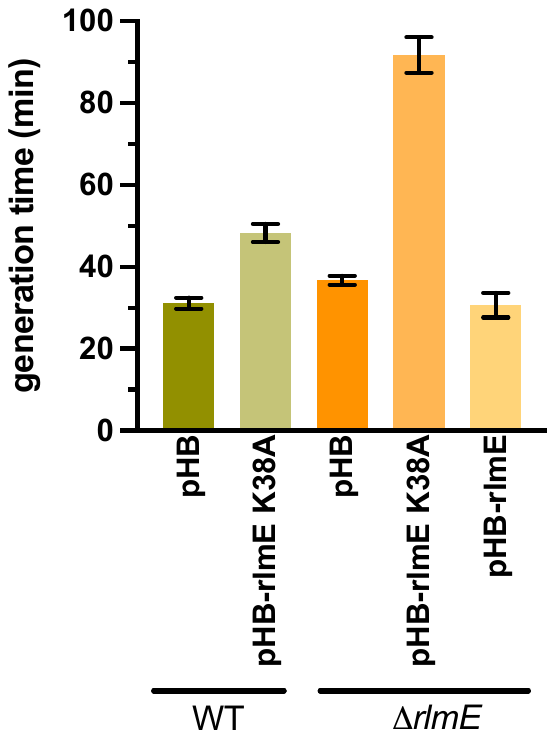

**Sup. Figure 2.** Rescue experiments by expressing functional or catalytically-inert RlmE enzymes in WT or *ΔrlmE* strains. The generation times were plotted. pHB is the low-copy plasmid backbone. Error bars are standard errors, n ≥ 7.

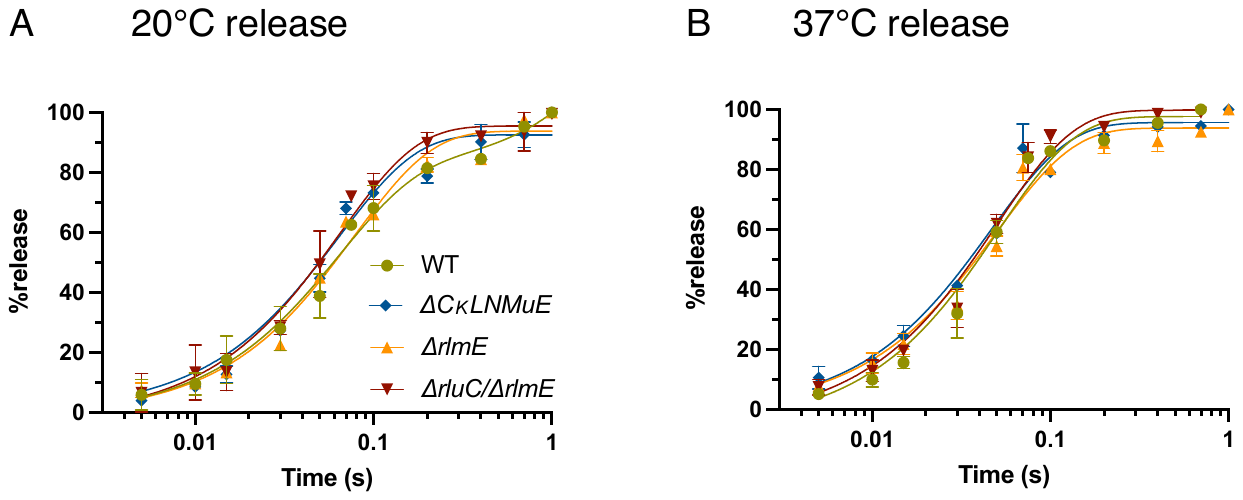

**Sup. Figure 3.** *In vitro* fast kinetics-based release assays of KO ribosomes. Time courses at 20℃ (A) and 37℃ (B). Error bars are standard errors.

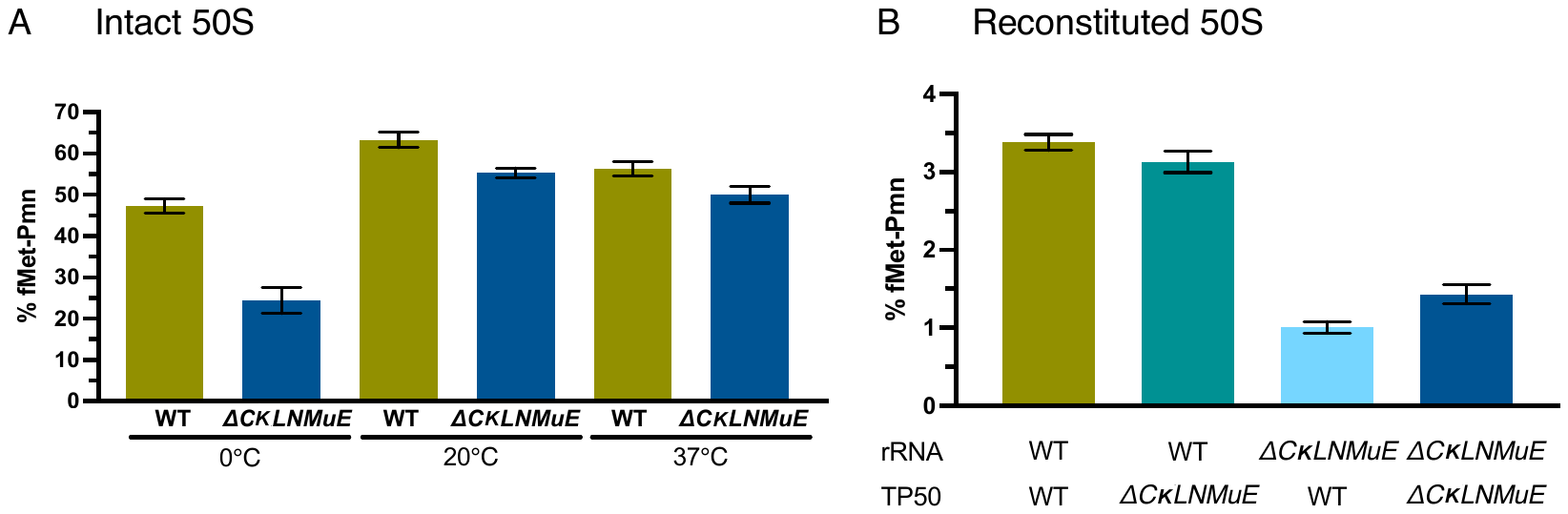

**Sup. Figure 4.** Comparison of fragment reactions at different temperatures. (A) Intact WT and *ΔCKLNMuE* 50S-catalyzed fragment reactions under standard conditions (on ice) or at higher temperatures. (B) Pairwise reconstituted 50S-catalyzed fragment reactions on ice for 20 min. Controls without 50S gave negligible signal. Error bars are standard errors, n ≥ 2.

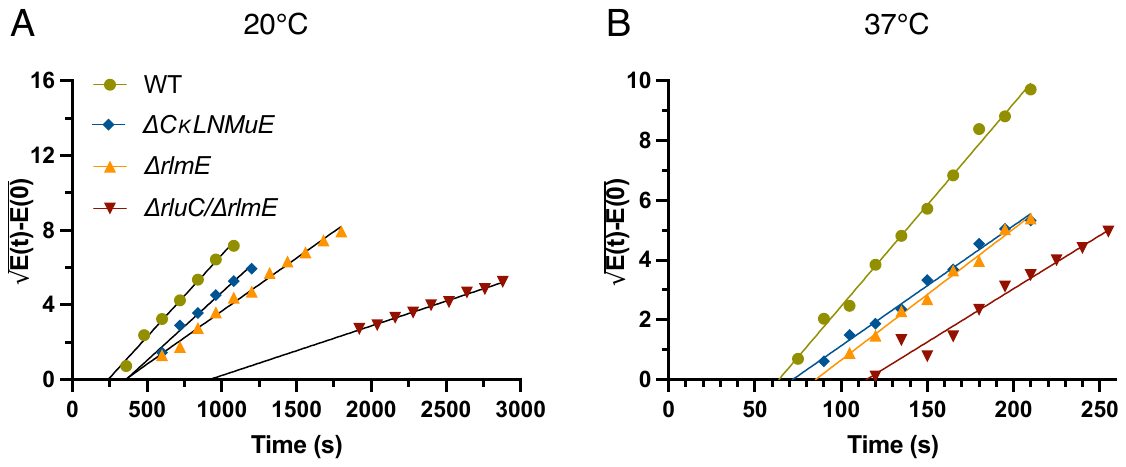

**Sup. Figure 5.** Time courses of β-galactosidase induction *in vivo* (Schleif *et al.*, 1973) at 20℃ (A) and 37℃ (B). E = enzyme activity. The X-axis intercepts of the linear parts of the Schleif plots indicated the time (T_first_) for the ribosome to synthesize the first LacZ protein (1024 aa), so the elongation rates (see Fig. 5) were calculated as 1024/T_first_.

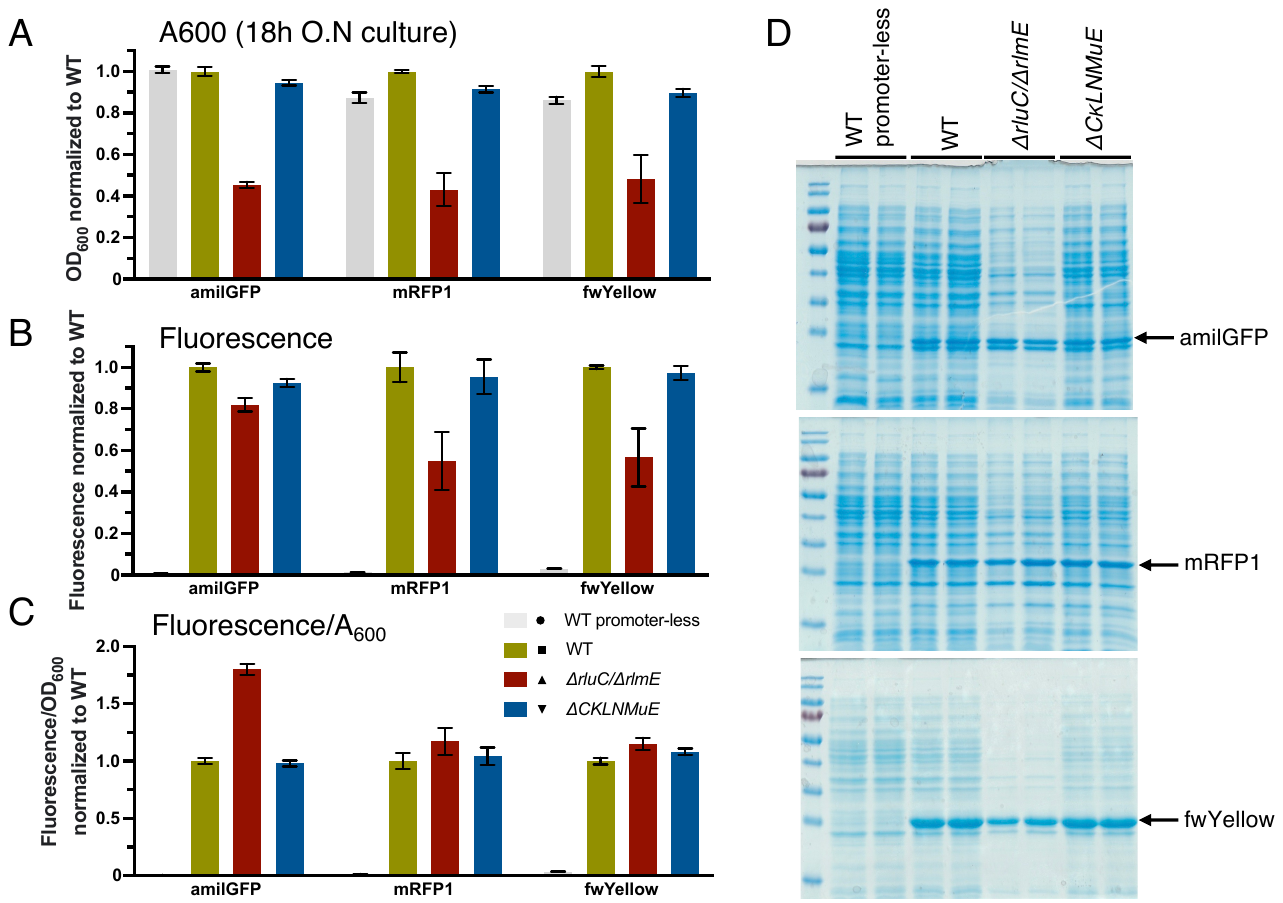

**Sup. Figure 6.** Influence of rRNA modification enzyme KOs on overexpression of fluorescent proteins and on growth. (A) Growth of the respective strains at 18h. (B) Constitutive overexpression at 37℃ of functional green (amilGFP), red (mRFP1) and yellow (fwYellow) fluorescent proteins encoded on high-copy plasmids. (C) Normalized fluorescence per cell. Error bars are standard errors, n = 6. (D) Gel analysis of influence of rRNA modification enzyme KOs on the total yield of overexpressed amilGFP, mRFP1 and fwYellow. Representative SDS-PAGE gels after 18h incubation at 37℃ show both functional and non-functional constitutively-expressed Coomassie-stained proteins. Arrows indicate induction bands.

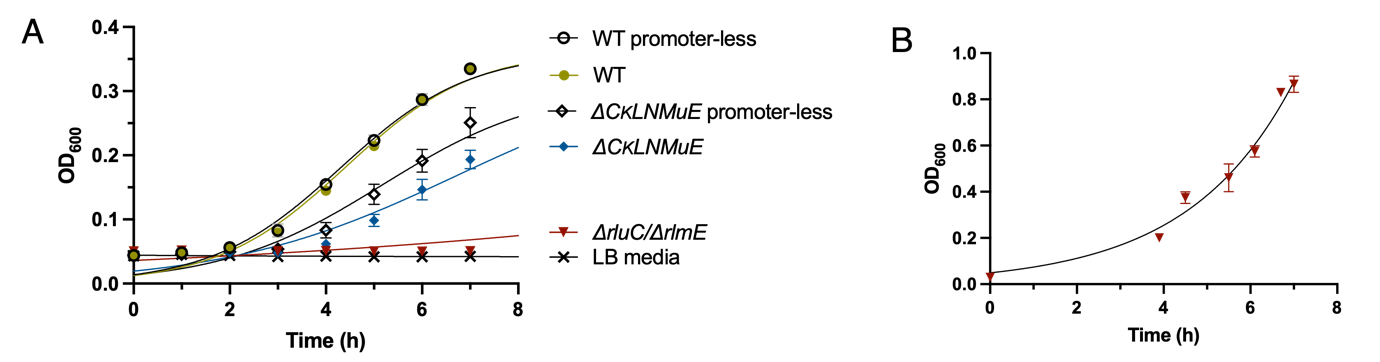

**Sup. Figure 7.** Influence of constitutive overexpression of mRFP1 on growth of *ΔCKLNMuE* and *ΔrlmE/Δrlu*C. (A) Growth of two combined KO strains when harboring mRFP1 encoded on a high-copy plasmid. In two controls, the plasmid lacked the promoter for mRFP1 (promoter-less). (B) Growth of *ΔrlmE/Δrlu*C without plasmid at 37℃. Different OD ranges were due to measurement with plate reader or spectrophotometer. Error bars are standard errors, n ≥ 3.
